# Externally provided rewards increase internal preference, but not as much as preferred ones without extrinsic rewards

**DOI:** 10.1101/2023.05.03.539192

**Authors:** Jianhong Zhu, Kentaro Katahira, Makoto Hirakawa, Takashi Nakao

## Abstract

It is well known that preferences are formed through choices, known as choice-induced preference change (CIPC). However, whether value learned through externally provided rewards influences the preferences formed through CIPC remains unclear. To address this issue, we used tasks for decision-making guided by reward provided by the external environment (externally guided decision-making; EDM) and for decision-making guided by one’s internal preference (internally guided decision-making; IDM). In the IDM task, we presented stimuli with learned value in the EDM and novel stimuli to examine whether the value in the EDM affects preferences. Stimuli reinforced by rewards given in the EDM were reflected in the IDM’s initial preference and further increased through CIPC in the IDM. However, such stimuli were not as strongly preferred as the most preferred novel stimulus in the IDM, indicating the superiority of intrinsically learned values (SIV). The underlying process of this phenomenon is discussed in terms of the fundamental self-hypothesis.

**Author Summary:** We make decisions based on internal value criteria, which are individual preferences, or based on external value criteria, which are the values learned from the external environment. Although it is known that values are learned in both types of decisions, is there a difference in the nature of these values? Our study uses simulation and fits human behavioral data to address this question. The results showed that stimuli that were learned to be highly valued because of external feedback became preferred in subsequent preference judgments. However, it is interesting to note that such stimuli were not chosen as much as stimuli that were preferred without influence from the external environment. This finding suggests that values formed through one’s own criteria have characteristics distinct from those formed through external environmental influence. Our findings promote an integrated understanding of the decision-making process.

## Introduction

Existentialism, according to Jean-Paul Sartre ^[1]^, supposed that the reason for one’s existence is not predetermined and that a sense of self is established through one’s own choices under the freedom given to one, as expressed in the phrase “Existence precedes essence.” In contrast, structuralism, which was developed by Claude Lévi-Strauss ^[2,3]^, emphasizes the influences on human behavior from the external environment, such as social and cultural structures. In everyday life, after making a decision, the feedback provided by the external environment reflects the value criteria of the external environment. However, do external value criteria influence decision-making when people can rely solely on their internal values? Moreover, is there any difference between the values formed by the choices one makes based on one’s internal criteria and the values produced by the external environment?

Depending on the situation, we make decisions based on criteria given by the external environment (such as a monetary reward), known as externally guided decision-making (EDM), or on one’s own internal criteria (such as a sense of value, beliefs, and preferences), referred to as internally guided decision-making (IDM). Although EDM and IDM are distinct in their conceptual definition, experimental operation, and neural bases ^[4,5,6,7,8]^, they are similar in that the option’s values are learned through decision making ^[6,7,9,10,11]^.

In EDM, the value of an option is considered to be updated based on the predicted value and actual feedback given after the decision making ^[12,13,14,15,16,17,18]^. For example, in a reward learning task where one item is chosen from two items and each is rewarded with a certain probability, the item’s value increases when rewarded feedback is received ^[12,18]^.

In IDM tasks, such as a preference judgment task, no externally delivered feedback indicates a correct answer. Even in such a case, an item’s value (preference) changes with the choice itself, not based on the feedback ^[19,20,21]^. More specifically, in the preference judgment task of choosing a preferred item from two presented items, the value of the chosen item increases while the value of the rejected item decreases (choice-induced preference change, CIPC ^[9,19]^).

EDM studies have progressed in mathematical understanding by analyzing behavioral data using computational models that represent the information process ^[22,23,24,25]^. The model of reinforcement learning (RL) explains the value learning mechanism in EDM. In the standard RL model, the choice behavior is guided by the expected value (e.g., $0.80 is the expected value of $1.00 dollar being compensated with an 80 percent probability) associated with the option. The expected value is updated according to the prediction error (i.e., the difference between the provided reward and the expected value) ^[22,26,27,28]^. The suitability of the computational model has been tested by fitting the model to the trial-by-trial choice of behavioral data. The model-based analysis can be used to estimate model parameters, such as learning rate (the degree of value update), and latent variables, such as the expected value and prediction error of each trial. Furthermore, these estimated parameters and latent variables have been used in neuroscience to explain the neural substrates of EDM _[19,29,30]._

For IDM, CIPC has been examined for many years using changes in subjective preference ratings as an index ^[21,31,32,33]^. In recent years, choice-based learning (CBL) models have been proposed ^[6,7,10,11]^, and computational modeling has progressed. CBL models are based on the RL model, but unlike RL, they use choice behavior itself instead of feedback from the external environment. In addition, differing from the typical RL, a CBL model was verified that updates the value of both chosen and rejected items ^[9]^. Thus, in the CBL model, chosen items are treated as correct answers, while rejected items are treated as incorrect answers to update their value. Although the EDM and IDM differ in the kinds of choice results they have, they update values based on feedback from the external environment or their own choice. Therefore, they are similar in that they update values based on the difference between expected values and the results of choice.

Since EDM and IDM have been studied as independent research areas, there is much ambiguity about their relationship. Although comparative studies of EDM and IDM have been reported in recent years, these studies have focused on their differences ^[4,5,6,7,8,34]^. Therefore, whether the value learned through the EDM affects the value of the IDM has not been explored. It has been reported that reward-related neural responses are involved in the value-learning process in IDM ^[6,21,31,35,36,37]^, as with EDM ^[38,39,40,41,42]^. From these findings, it can be inferred that the values learned through these two decision-making processes are not clearly distinct, and the values learned in EDM can have an impact on IDM. However, those studies examine EDM or IDM independently and the similarity of the neural basis provides insufficient evidence for the impact of values in EDM on IDM. Thus, it remains unclear whether the values learned in the EDM affect the IDM.

This study investigated whether, how, and to what extent the value learned in EDM affects the value in IDM. We used simple EDM and IDM tasks with novel contour shapes; the IDM task followed the EDM task. In the IDM task, the same stimuli used with the EDM task were used for the four (two high and two low reward probability) stimuli in addition to eleven novel stimuli.

We first tested whether the values learned in EDM affect IDM from classical model-free behavioral data analysis using the chosen frequency of items in IDM ^[5,7]^. If the value learned in the EDM affects the IDM, it is predicted that stimuli with higher values learned in the EDM will be chosen more frequently than novel stimuli in the IDM. It is also expected that stimuli with lower values learned in the EDM will be chosen less frequently than novel stimuli in the IDM.

To examine how the values in EDM reflected IDM, we applied four computational models (see Table 1 in Methods) to the IDM data, compared the models, and investigated the estimated initial values of each stimulus in the IDM. The initial values of different stimulus types in the IDM in these models differed (Table 1). Model 1, in which the initial values of all stimuli were the same, indicated that the values learned in the EDM did not affect the IDM. Model 2, in which only the initial value of the high-reward probability stimulus differed from the other stimuli, indicated that the high value learned in the EDM affected the IDM. Model 3, in which only the initial value of the low-reward probability stimuli differed from the other stimuli, shows that the low value learned in the EDM affects the IDM. Model 4, in which the initial values of all types of stimuli were different, illustrates that high and low values learned in the EDM affect the IDM. If the value in the EDM is reflected in the initial value in the IDM, we predict that other models will fit behavioral data better than Model 1. In addition, if both high and low values in the EDM were reflected in the IDM, then Model 4 would be the best fit for this behavior. For the computational model, simulations were conducted to confirm that the model parameters could be adequately estimated (parameter recovery) and that an appropriate model could be selected (model recovery) before the behavioral data analyses.

**Table 1.** The settings of initial values of each stimulus type in different CBL models.

| Model | Novel | HP | LP |
| --- | --- | --- | --- |
| Model 1 | | $\eta$ | |
| Model 2 | 0.5 | $\eta_{HP}$ | 0.5 |
| Model 3 | 0.5 | 0.5 | $\eta_{LP}$ |
| Model 4 | 0.5 | $\eta_{HP}$ | $\eta_{LP}$ |

Note: $\eta$ denotes the free parameter for initial value in IDM. HP and LP denote high- probability reward stimuli (90%, 80%) and low probability reward stimuli (20%, 10%) in the EDM task, respectively.

To further investigate how the value learned in EDM affects IDM, we explored whether the values learned in EDM affect the degree of value change in IDM. To examine the value changes from initial to final, we used values estimated by models that best fitted the behavioral data. Stimuli that reflect high or low values in the EDM are expected to be chosen or rejected more frequently in the IDM, respectively, thereby further changing their values. As all four models described above with different initial value settings assumed value changes in the IDM based on a previous study ^[9]^, an additional computational model analysis was performed to rule out the possibility that the value did not change in the IDM.

Finally, by synthesizing the effects of the EDM on the initial value and the value change in the IDM, the effect of the EDM on the IDM was examined. To this end, we compared the final values of the IDM between the EDM stimuli and all the novel stimuli. If the influence of the value in the EDM is dominant, then it is expected that stimuli with high values in the EDM will ultimately be the highest-value stimuli in the IDM. Similarly, a stimulus with a low value in the EDM is expected to have the lowest final value in the IDM. On the contrary, the value learned solely in IDM may have a different specificity compared to the value influenced by the external environment. In this case, a novel stimulus in IDM with a higher value could eventually become more valuable than a stimulus with a higher value in EDM. To confirm the validity of the final values estimated by the computational model, we examined the consistency between the final values estimated by the model and the chosen frequency of each stimulus in the IDM or the subjective ratings of each stimulus after the IDM task.

We confirmed whether the participants successfully learned the value of each stimulus in the EDM from the correct response rate. Therefore, in this study, a computational model analysis of the EDM behavioral data is not necessary. For reference, we report the results of the RL model analysis for EDM in the supplementary materials.

## Results

### Correct response rate in EDM task

We confirmed that participants learned the value of stimuli through EDM by investigating the results of the correct response rate (Fig 1a). The correct response rate of the 80% vs. 20% stimulus pair was 0.698 (*SD* = .093), and the 90% vs. 10% stimulus pair was 0.830 (*SD* = .092), both of which were higher than chance level (0.5) (*t*s(37) > 13.124, *p*s < .001, 95% CI = 0.667, 0.728 in 80% vs. 20% stimulus pair, 95% CI = 0.800, 0.861 in 90% vs. 10% stimulus pair). In addition, there was also a significant difference between these two conditions (*t*(37) = 8.189, *p* < .001, *d* = 1.434, 95% CI = 0.100, 0.165).

**Fig 1.**
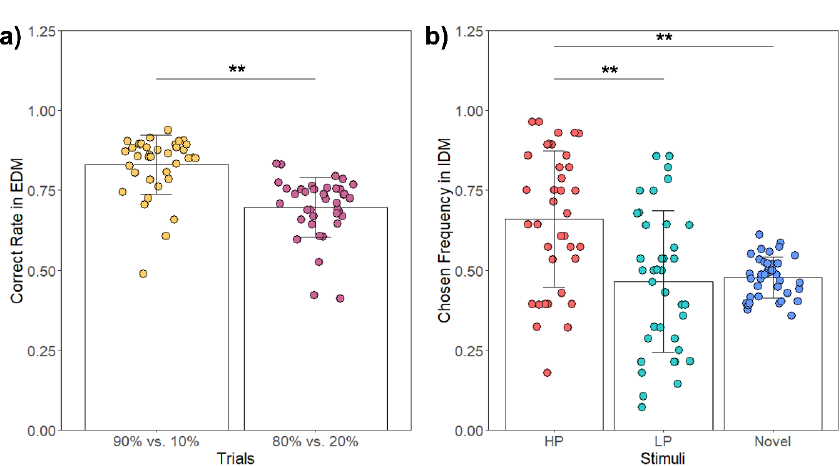
Results of model-free behavioral data indicators. a) Mean correct response rate in the EDM task. The easier condition was stimulus pairs with 90% vs. 10% probability of obtaining a reward, and the harder condition was stimulus pairs with 80% vs. 20% probability of obtaining a reward. b) Mean chosen frequency of each stimulus type in the IDM task. The error bars and colored dots of all figures indicated *SD* and each participant’s data, respectively. ** *p* < .001.

### Chosen frequency in IDM task

To examine whether the values learned in EDM affect IDM, we examined the chosen frequency of each stimulus type in IDM (Fig 1b). In the IDM task, there was no difference in the frequency of choice between the 90% (chosen frequency = 0.675) and 80% (chosen frequency = 0.645) stimuli presented in EDM (*t*(37) = 0.552, Holm-adjusted *p* = 1.000, *d* = 0.110, 95% CI = −0.080, 0.140), or between the 10% (chosen frequency = 0.445) and 20% (chosen frequency = 0.485) stimuli presented in EDM (*t*(37) = −0.601, Holm-adjusted *p* = 1.000, *d* = −0.130, 95% CI = −0.173, 0.094). Therefore, 90% and 80% stimuli were grouped together as high-probability (HP) stimuli, and 10% and 20% stimuli as low-probability (LP) stimuli.

The mean chosen frequency and standard deviation (*SD*) of the three types of stimuli were 0.660 (0.213), 0.465 (0.222), and 0.477 (0.064) for HP, LP, and novel stimuli, respectively. Participants preferred to choose HP stimuli as the preferred item than novel stimuli and LP stimuli (*t*s(37) > 4.211, Holm-adjusted *p*s < .001, *d*s > 0.886). However, there was no significant difference between their preference for the LP and novel stimuli (*t*(37) = −0.268, Holm-adjusted *p* = .790, *d* = −0.073, 95% CI = −0.518, 0.372). These results showed that only the high value of stimuli learned in the EDM task influenced the IDM task.

### Simulation 1 (Parameter recovery)

Fig 2 shows the results of parameter recovery for each of the four CBL models (Table 1). We confirmed strong consistency between the set parameter values (simulated) to generate artificial behavioral data and the estimated values (fitted) by fitting the model generating the data in all CBL models (*r*s > .604).

**Fig 2.**
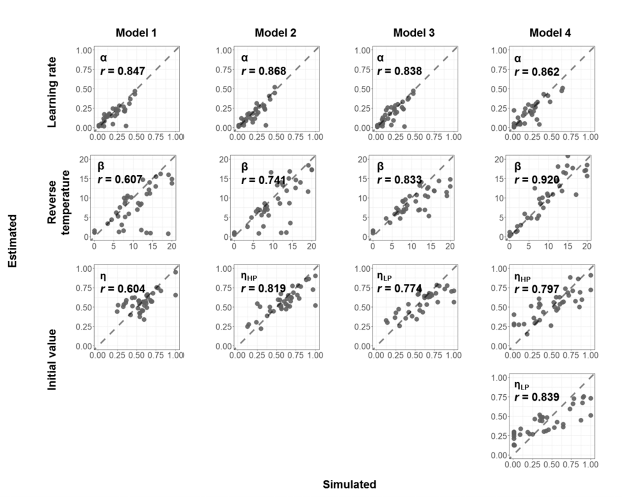
Results of parameter recovery simulation. This simulation was conducted to confirm whether each model could be well estimated as the set value of each parameter. The correlation coefficient between simulated and fitted was shown as parameter recovery indices.

### Simulation 2 (Model recovery)

Table 2 presents the widely applicable Bayesian information criterion (WBIC) estimated when data generated by the individual models were fitted to all models. In the comparison of the WBICs, the smaller the value of the WBIC, the higher the fit of the generated data to the model. As shown in Table 2, all models could complete model recovery well. Whichever model generated the data had the highest match to the same model.

**Table 2.** Results of WBIC for model recovery for Model 1–4 shown in Table 1.

| Simulated \ Fitted | Model 1 | Model 2 | Model 3 | Model 4 |
| --- | --- | --- | --- | --- |
| Model 1 | **2135.055** | 2144.160 | 2136.063 | 2150.349 |
| Model 2 | 2100.200 | **2082.634** | 2107.459 | 2093.550 |
| Model 3 | 2113.086 | 2118.352 | **2089.181** | 2099.140 |
| Model 4 | 2103.990 | 2077.184 | 2090.915 | **2073.248** |

Note: Bold numbers in the table represent the WBIC values of the models that best fit the artificial data.

We calculated the Bayes factor (BF) to compare which model had the higher probability of generating data. When Model X was the true model (i.e., the model used to generate artificial data), the result of the BF between the Model X and Y was shown by BF*_XY_*. The marginal likelihood of Model X and Y was the numerator and denominator, respectively. When Model 1 was the true model, the evidence was not as strong for Model 1 when compared to Model 3 (BF*_13_* = 2.740), indicating that the probability of data generated by Model 1 was 2.740 times that of Model 3. However, there was very strong evidence for Model 1 when compared to Models 2 and 4 (BF*_12_* = 9000.181, BF*_14_* = 4.386×10^6^). When Models 2 and 3 were the true models, there were very strong supports for each model compared to the other models (BF*_21_* = 4.254×10^7^, BF*_23_*= 6.045×10^10^, BF*_24_* = 5.505×10^4^, BF*_31_* = 2.409×10^10^, BF*_32_* = 4.664×10^12^, BF*_34_* = 2.114×10^4^). When Model 4 was the true model, strong evidence was found for comparison with Model 2 (BF*_42_* = 51.213), and very strong evidence for comparison with Models 1 and 3 (BF*_41_*= 2.244×10^13^, BF*_43_* = 4.706×10^7^).

### Model fit to behavioral data

To examine how the EDM values reflect the IDM, we fit the four models to actual IDM behavioral data. The results shown in Table 3 indicated that Model 2 had a better fit than other models. Model 2 was a model in which only the value of the HP stimuli in EDM was reflected in the initial value in IDM. This result was consistent with the result of chosen frequency in Fig 1b. We calculated BF for inter-model comparison between Model 2 and other models. The BF was calculated using the marginal likelihood of Model 2 as the numerator. The results showed that there was strong evidence to support Model 2 (BF*_21_* = 3.302×10^6^, BF*_23_* = 2.913×10^8^, and BF*_24_* = 7.922×10^4^).

**Table 3.** WBIC results of IDM behavioral data fit with Model 1–4 shown in Table 1.

| Model | WBIC |
| --- | --- |
| Model 1 | 2189.52 |
| **Model 2** | **2174.51** |
| Model 3 | 2194.00 |
| Model 4 | 2185.79 |

Note: Bold numbers in the table represent the WBIC values of the model that best fit the behavioral data.

In addition, to estimate the extent to which Model 2 explained behavior, we compared it to a model with random choice that did not predict behavior at all (see the supplementary materials for more details). The BF results showed that Model 2 was 4.130×10^256^ times more likely to explain behavior than the random model.

### Confirmation of estimated initial values in IDM

The mean initial value of the HP stimuli in the IDM estimated using the CBL model (Model 2) was 0.575 (Fig 3a). To confirm the values learned in EDM were reflected in the initial values in IDM, we compared the estimated initial value for HP, fixed initial values (0.5) for LP, and novel stimuli. We found that HP stimuli had a higher initial value than LP stimuli and novel stimuli (*t*(37) = 7.057, *p* < .001, 95% CI = 0.554, 0.597). These results showed that the high value learned in EDM was reflected in the initial value in IDM.

**Fig 3.**
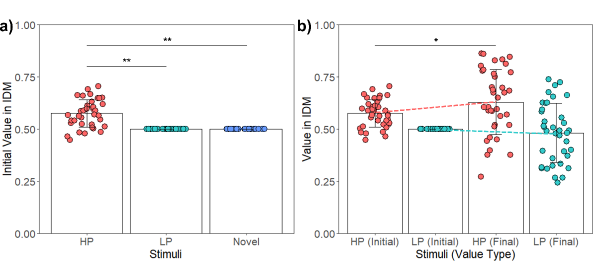
Values for stimuli estimated by the computational models. a) Mean initial value of each stimulus type estimated by Model 2 in IDM task. b) Mean initial and final values for HP and LP stimuli in the IDM task. Dotted lines represent the value change trend from initial to final of each stimulus type. The red and green represent the change trend of HP and LP stimuli, respectively. The error bars and colored dots of all figures indicate *SD* and each participant’s data, respectively. * *p* < .05, ** *p* < .001.

### Estimated value change from initial to final values of HP and LP stimuli in IDM

To examine whether the value learned in EDM affected the degree of value change in IDM, we compared the changes in value between HP and LP stimuli (Fig 3b). We performed a two-factor repeated measures ANOVA for the stimuli type (HP stimuli and LP stimuli) and the value type (initial value and final value), which found no significant main effect in the value type (*F*(1, 37) = 1.307, *p* = .260, partial *η^2^* = .034), but a significant main effect for the stimuli type (*F*(1, 37) = 37.072, *p* < .001, partial *η^2^* = .500), and the interaction (*F*(1, 37) = 10.500, *p* = .003, partial *η^2^* = .221). We compared initial value and final value in each stimulus type, and found the final values were higher than the initial values for HP stimuli (*t*(37) = −2.408, Holm-adjusted *p* = .021, *d* = −0.582, 95% CI = −0.101, −0.009), whereas no difference was found for LP stimuli (*t*(37) = .791, Holm-adjusted *p* = .434, *d* = 0.142, 95% CI = −0.040, 0.070).

Thus, results showed that the HP stimuli had a higher initial value in IDM due to the influence of EDM and therefore was chosen frequently in IDM, making it increase in value in IDM.

### Additional model comparison between the Model 2 and the models without value updates in IDM

Although we observed that HP stimuli were further valued in IDM based on computational model analysis, Model 2 used in the analysis assumes that the values change. The validity of this assumption was not examined in the comparisons among the four models with different initial value settings (Table 1). Therefore, an additional computational model analysis was performed to investigate whether the value did not change in the IDM. Additional models without value changes were constructed based on Model 2: Specifically, we created four additional models (Table 4): Model A, in which only the value of the HP stimuli was not updated; Model B, in which both the HP and LP stimuli were not updated and only the value of the novel stimuli was updated; Model C, in which only the value of the novel stimuli was not updated; and Model D, in which the values of all the stimuli were not updated. All new models were analyzed through simulations and the fitting of behavioral data.

**Table 4.**
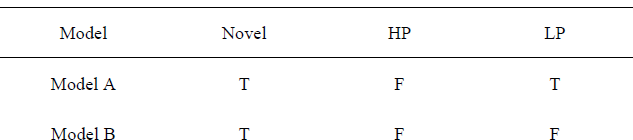
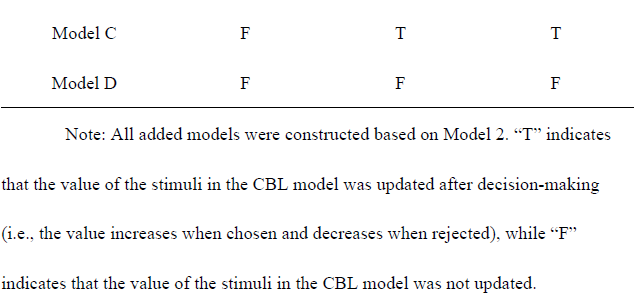
The settings of value updating of each stimulus type in the added models (A–D) without value updates in IDM.

| Model | Novel | HP | LP |
| --- | --- | --- | --- |
| Model A | T | F | T |
| Model B | T | F | F |
| Model C | F | T | T |
| Model D | F | F | F |

Note: All added models were constructed based on Model 2. “T” indicates that the value of the stimuli in the CBL model was updated after decision-making (i.e., the value increases when chosen and decreases when rejected), while “F” indicates that the value of the stimuli in the CBL model was not updated.

The results of the parameter recovery for all added models are shown in Fig 4. Excluding Model D, there was consistency between the set parameter values (simulated) and the estimated values (fitted) for the remaining three added models (*r*s > .52).

**Fig 4.**
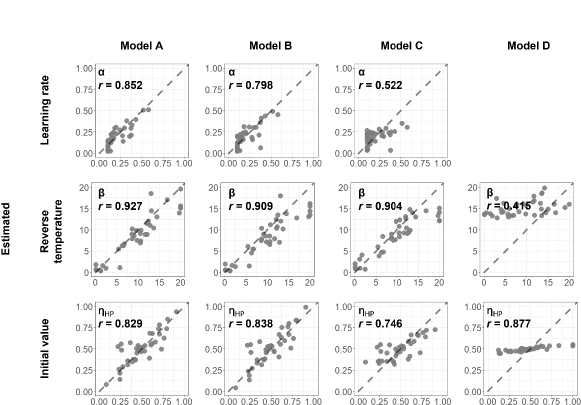
Results of parameter recovery simulation for additional comparison between the Model 2 and the added models (A–D) without value updates in IDM. This simulation was conducted to confirm whether each model could be well estimated as the set value of each parameter. The correlation coefficient between simulated and fitted was shown as parameter recovery indices.

Model recovery was confirmed for all models except Model D (see Table 5). When the same true model (used to generate artificial data) was used for the analysis, the fit of the analytical model to the artificial data was the best. When Model 2 and Models A–C were the true models, strong evidence was found for comparison with other models (BF*_2A_*= 1.299×10^38^, BF*_2B_* = 1.764×10^79^, BF*_2C_* = 2.399×10^135^, BF*_2D_* = 2.264×10^184^, BF*_A2_* = 1.176×10^9^, BF*_AB_* = 1.564×10^35^, BF*_AC_*= 2.425×10^144^, BF*_AD_* = 2.885×10^158^, BF*_B2_* = 5.043×10^17^, BF*_BA_* = 9.821×10^11^, BF*_BC_* = 5.411×10^141^, BF*_BD_* = 6.590×10^141^, BF*_C2_* = 1.695×10^48^, BF*_CA_* = 3.551×10^73^, BF*_CB_* = 1.915×10^84^, BF*_CD_* = 2.680×10^79^). When Model D was the true model, although positive evidence was found for comparison with Models 2, A, and B (BF*_D2_* = 12.718, BF*_DA_* = 11.179, and BF*_DB_* = 3.251), no evidence was found for comparison with Model C (BF*_DC_* = 0.139).

**Table 5.** Results of WBIC for model recovery of additional comparison between Model 2 and the added models (A–D) shown in Table 4.

| Fitted<br>Simulated | Model 2 | Model A | Model B | Model C | Model D |
| --- | --- | --- | --- | --- | --- |
| Model 2 | **2342.541** | 2430.301 | 2525.013 | 2654.265 | 2767.034 |
| Model A | 2427.446 | **2406.561** | 2487.599 | 2739.019 | 2771.429 |
| Model B | 2486.038 | 2472.889 | **2445.276** | 2771.629 | 2771.826 |
| Model C | 2691.428 | 2749.732 | 2774.443 | **2580.376** | 2763.266 |
| Model D | 2772.947 | 2772.818 | 2771.583 | **2768.434** | 2770.404 |

Note: Bold numbers in the table represent the WBIC values of the models that best fit the artificial data.

Although Model D did not adequately complete the simulations, we compared all the models by fitting them to the behavioral data. The results indicate that Model 2 had a better fit than the other models. (Table 6; BF*_2A_* = 2.424×10^47^, BF*_2B_* = 1.106×10^107^, BF*_2C_* = 2.433×10^147^, BF*_2D_* = 1.345×10^232^).

**Table 6.** WBIC results of IDM behavioral data fit with Model 2 and added models.

| Model | WBIC |
| --- | --- |
| **Model 2** | **2170.18** |
| Model A | 2279.29 |
| Model B | 2416.66 |
| Model C | 2509.55 |
| Model D | 2704.68 |

Note: Bold numbers in the table represent the WBIC values of the model that best fit the behavioral data.

These results confirmed that the values of all stimuli including HP in the IDM were updated after the choice.

### Estimated final values in IDM

To examine the effect of the value learned in the EDM on the IDM, we compared the final value of the HP and LP with each novel stimulus in IDM. We used the best-fit model (Model 2) to estimate the values. We ranked all novel stimuli in order of their final value from high to low within each participant (those labeled as N1 to N11) and then compared them with HP or LP stimuli (Fig 5a). The results showed that HP stimuli were lower than N1 (*t*(37) = −2.883, Holm-adjusted *p* = .026, *d* = − 0.744, 95% CI = −0.157, −0.027) and higher than N5 to N11 (*t*s(37) > 3.133, Holm-adjusted *p*s < .05, *d*s > 0.854). LP stimuli were lower than N1 to N4 (*t*s(37) < −2.768, Holm-adjusted *p*s < .05, *d*s < −0.742) and higher than N8 to N11 (*t*s(37) > 2.650, Holm-adjusted *p*s < .05, *d*s > 0.749).

**Fig 5.**
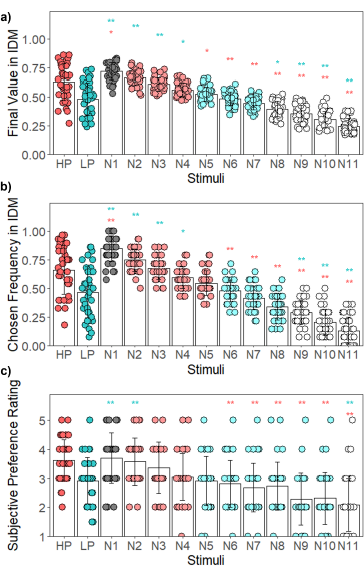
Comparison of the final value estimated by the Model 2 among stimulus type (a) and the correspondence between that estimated final value and chosen frequency (b) or the subjective preference ratings (c) a) Comparison of the mean final value of HP and LP stimuli in IDM task with all novel stimuli. N1–N11 denote the rank of the novel stimuli in the final value. N1 was the most favorite novel stimulus, and N11 was the least favorite novel stimulus for each participant. The red “*” represents the comparison result with HP, and the green “*” represents the comparison result with LP. There was no difference between N2–N4 and HP and between N5–N7 and LP. The error bars and colored dots indicate *SD* and each participant’s data, respectively. * *p* < .05, ** *p* < .001. b) Comparison of the mean chosen frequency in IDM for all types of stimuli. N1–N11 denote the rank of the novel stimuli in the final value. The error bars and colored dots indicate SD and each participant’s data, respectively. * *p* < .05, ** *p* < .001. Comparison of the subjective preferences ratings rated on a 5-point Likert scale (1 = Extremely Dislike, 5 = Extremely Like). N1–N11 denote the rank of the novel stimuli in the final value. The error bars and colored dots indicate *SD* and each participant’s data, respectively. * *p* < .05, ** *p* < .001.

Moreover, we compared the HP and LP with the mean value of all novel stimuli. Results showed that the final value of HP stimuli was higher than that of novel stimuli (*t*(37) = 4.708, Holm-adjusted *p* < .001, *d* = 1.254, 95% CI = 0.083, 0.207). In contrast, LP stimuli did not differ from novel stimuli (*t*(37) = −0.091, Holm-adjusted *p* = .928, *d* = −0.025, 95% CI = −0.060, 0.055).

These results showed that HP stimuli maintained their high value through the end of IDM, while their value was lower than the most preferred novel stimuli. That is, although the IDM is affected by the EDM value, the superiority of intrinsically learned values (SIV) was concurrently observed in the IDM. Regarding LP stimuli, there was no significant effect on the IDM final value as in the results for initial values, and LP stimuli were the average position of the novel stimuli.

### Consistency between the final value estimated by Model 2 and the chosen frequency or subjective preference ratings

To confirm the validity of the model-estimated final values, we examined the consistency between the model-estimated final values and the chosen frequency of each stimulus in the IDM or the subjective ratings of each stimulus after the IDM task. Comparisons were made between novel stimuli ranked by the final value (N1– N11), HP and LP stimuli for each chosen frequency, and subjective preference ratings. Fig 5b shows the consistency between the final value and chosen frequency. The results demonstrated that the chosen frequency of the HP stimuli was lower than that of the N1 stimulus (*t*(37) = −4.267, Holm-adjusted *p* < .001, *d* = −1.083, 95% CI = −1.561, −0.605), but higher than that of the N6–N11 stimuli (*t*s(37) > 3.905, Holm-adjusted *p*s < .01, *d*s > 1.083). The LP stimuli were lower than the N1–N4 stimuli (*t*s(37) < −2.783, Holm-adjusted *p*s < .05, *d*s < −0.747) but higher than the N9–N11 stimuli (*t*s(37) > 3.772, Holm-adjusted *p*s < .01, *d*s > 1.025).

Fig 5c depicts the consistency between the final value and the subjective preference ratings. The results showed that the subjective preference for the N1 stimulus with the highest final value was higher than that for the N4–N11 stimuli (*t*s(37) > 4.197, Holm-adjusted *p*s < .01, *d*s > 0.779). The subjective preference of the HP stimuli with the second highest final value was higher than that of the N5–N11 stimuli (*t*s(37) > 3.634, Holm-adjusted *p*s < .05, *d*s > 0.880), while the subjective preference of the LP stimulus was lower than that of the N1–N2 (*t*s(37) < −4.026, Holm-adjusted *p*s < .01, *d*s < −1.263), but higher than that of the N11 (*t*(37) = 4.295, Holm-adjusted *p* < .01, *d* = 0.971, 95% CI = 0.499, 1.443).

Overall, the results validated the final value estimated by Model 2, indicating that stimuli with high values in the IDM had a higher chosen frequency and were subjectively preferred.

## Discussion

The goal of this study was to determine whether, how, and to what extent the EDM value affects the IDM. Through a comparison of the chosen frequency of the three types of stimuli (novel, HP, LP) in the IDM task, the chosen frequency for HP stimuli was higher than LP stimuli and novel stimuli (Fig 1b). These results indicated that the values learned in EDM affect the IDM.

We conducted a computational model analysis to examine how the EDM values reflect the IDM. The model comparison revealed that Model 2, in which the initial values of the HP stimuli in the IDM were different from that of the other stimuli, best fit the data (Table 3). By comparing the initial values of each stimulus estimated by this model, we confirmed that the EDM values of the HP stimuli were reflected in the initial IDM values (Fig 3a). This result was consistent with the results of chosen frequency (Fig 1b). Collectively, the high values learned in EDM (reward learning task) were reflected in the initial values of IDM (preference judgment). This suggests a close relationship between the values obtained using the EDM and IDM.

One might argue that this finding reflects previous results from the extinction procedure; subjects in conditioning studies continued to choose the reinforced option even after removing the reward ^[43,44,45,46,47]^. However, in such cases, unlike the present study, the subjects were not explicitly told that the decision task or situation had changed. They were also placed in a position in which they expected that choosing the reinforced option would eventually reward them ^[43,44]^. In contrast, participants in the present study were clearly instructed on the difference between the EDM and IDM, and they were aware that they would not be rewarded in the IDM. That is, the present study operationally eliminated the participants’ choice of reinforced items based on the expectation of an externally derived reward and asked them to choose a preferred shape according to their own preferential criteria in the IDM. Furthermore, in the extinction procedure, the value of stimuli or behavior decreases when external rewards are not provided after the choice, and the option is gradually not chosen ^[45,46,47]^. In contrast, HP stimuli in the present study were more frequently chosen in the IDM, even though no reward feedback was presented, and their value further increased (Fig 3b). This difference suggests that, in the elimination procedure, the highly valued option continued to be chosen in anticipation of rewards from the external environment. In the IDM in this study however, high-value stimuli in the EDM were chosen based on their own preferences, resulting in a CIPC. Therefore, the results of the present study are not consistent with the sustained choice of highly rewarding stimuli reported in conditioning studies ^[45,46,47]^.

To further examine how the value learned in EDM affects IDM, we examined whether the value learned in EDM affects the degree of value change in IDM. The high value of HP at the end of IDM was not simply a result of the maintenance of the high initial value reflected as the effect of EDM, but it was also shown that the value of HP was further increased by selection in IDM (Fig 3b). Although we also examined the possibility that the value of HP stimuli was not updated in the IDM (Table 4), such a model did not fit well with the behavioral data (Table 6). Those results demonstrate that what is learned to be of high value according to externally derived criteria will subsequently be further valued within that individual through their own choices and the following CIPC. This result may indicate part of the internalization process of the value of the external environment if the values in the external and internal criteria differ.

To determine the extent to which the values learned in the EDM affected the IDM, we compared the model-estimated final values for all stimuli in the IDM. Interestingly, we found the superiority of intrinsically learned value (SIV) in the IDM, in which the most preferred novel stimulus learned in IDM (N1 in Fig 5a) was preferred over HP stimuli. If the values in the EDM and IDM were the same, then it would be expected that the stimuli with a higher value in the EDM would also have the highest value in HP in the final value in the IDM. This is because it was learned as highly valued in the EDM and subsequently chosen (Fig 1b) and valued in the IDM (Fig 3b). However, the SIV of novel stimuli is observed in the IDM (Fig 5a), indicating that our preferences are strongly influenced not only by externally given rewards but also by increased preferences on our own choices. Although the EDM value affects the IDM value, it is unlikely that the EDM and IDM values are identical.

However, the mechanisms underlying SIV in the IDM remain unclear. The recently proposed fundamental self-hypothesis ^[48,49,50,51,52,53,54]^ is considered relevant. This postulates that the self is a fundamental brain function that precedes and controls cognitive functions, such as perception, emotion, and reward, which has been proposed in studies of spontaneous brain activity ^[49,50,51]^ and the self-prioritization effect (SPE) ^[48,52,53,54]^. In this hypothesis, the self is embedded in spontaneous brain activity, and when a stimulus appears, the default mode network (DMN), which is responsible for processing self-associated stimuli, interacts with a task-related network to influence cognitive processing. A meta-analysis of the neural basis of IDM and EDM also confirmed that IDM differs from EDM in that the DMN is its primary neural substrate ^[4]^. In addition to the conceptual and operational differences between the EDM and IDM, there is a difference in task demands (i.e., whether decisions are made based on value criteria given by the environment or based on one’s own value criteria), that is, an essential difference in self-involvement. The continuous choice of stimuli as one’s own favorite shape, rather than because it has previously been rewarded, is likely to increase the self-relatedness of the item. As self-related stimuli are known to induce reward-related brain activity ^[55,56]^, an increase in self-relevance may trigger an internal reward response ^[6,21,31,35,36,37]^, leading to an increase in value. As a result, the most preferred novel stimulus learned in the IDM might have a higher value than HP stimuli in the IDM.

Although the model-free measure of chosen frequency also confirmed this SIV (Fig 5b), which was not reflected in the subjective preference ratings after IDM, subjective preference showed no significant difference between HP and N1 (Fig 5c). In the IDM task, 15 stimuli were presented in different combinations in each of the 105 trials. It is possible that this complex task setting prevented the subjective recognition of which stimuli were most favorable or chosen. It is necessary to reexamine whether this SIV can be observed in subjective preference ratings by experimenting with simpler task settings.

The low values learned in EDM did not affect IDM. In the EDM task, when an incorrect answer was chosen, feedback was displayed as 0 (simply not presented with a reward) rather than presented with a punishment. There was a possibility that the non-reward of LP stimuli in EDM did not affect participants’ preference for those stimuli, and therefore the initial value of LP stimuli in IDM was the same as for novel stimuli. Depending on whether the value is learned through reward acquisition or punishment avoidance in EDM, it is possible that IDM will reflect either high or low value in EDM.

Is it possible to interpret the results of this study based on familiarity? It is thought that the stimuli presented in the EDM (i.e., HP and LP) are processed as more familiar stimuli in the IDM than as novel stimuli. Although familiarity effects cannot be completely ruled out, they alone cannot explain the results of this study. If familiarity could explain the behavioral data in the IDM, we would expect to observe the same differences in the chosen frequency between the LP and novel stimuli as between the HP and novel stimuli. Additionally, if familiarity is an important factor, we would expect a model like Model 4 (where HP and LP are different from novel stimuli) to fit the behavioral data better. However, Model 2 was adopted, in which the initial value of only the HP stimuli was different from the other stimuli. This suggests that it is more plausible to assume the influence of value rather than that of familiarity. Conversely, a caveat is that the influence of familiarity cannot be completely ruled out, and the value of the LP was possibly higher because of the influence of familiarity than it would have been without it. To examine the influence of familiarity, it would be necessary to compare the LP with a stimulus without feedback that was presented for the same duration as the LP before the IDM.

This study showed for the first time the value of EDM affects IDM, and the SIV in IDM. Nevertheless, the present studies have several main limitations. First, the study showed that high value in EDM was reflected in IDM and low value was not. However, we should note that the results did not lead to any general conclusions about the relationship between EDM and IDM values but were a conclusion that depended on the task settings of this study. In this study, EDM used relatively easy reward probability settings such as 90% vs. 10% and 80% vs. 20%, where learning of value was easily established, to examine whether the value of EDM was reflected in IDM. Therefore, there were few opportunities to receive incorrect feedback after choosing LP stimuli and decreasing their values: the mean proportion of trials in which incorrect feedback was given after choosing the LP stimulus was 0.103, with *SD* of 0.113 (for comparison, the mean proportion of trials in which correct feedback was given after choosing HP stimuli was 0.734, with *SD* of 0.150). As a result, participants likely learned that LP stimuli were of relatively low value but did not come to the realization that they had to actively avoid LP stimuli. Therefore, it is assumed that LP stimuli in IDM were treated as having the same initial value as novel stimuli. When using an EDM task where the participant actively decides not to choose an item to avoid losses, the low value in EDM may affect IDM. Therefore, there is room for further study on this point.

Second, the possibility of explanations using other types of models has not yet been explored. For example, the cognitive dissonance theory has been used to explain the phenomenon of CIPC in IDM ^[57]^. In this theory, CIPC is explained by which choosing one item from two items with the same subjective preference rating, causing dissonant feelings (i.e., cognitive dissonance). Subsequently, they adjust their preferences for the chosen and rejected items in order to reinforce that their choice is reasonable. A difference between the cognitive dissonance theory and the CBL model is that the former assumes that preferences change in situations where one of the pairs of equal liking is chosen. The latter assumes that preferences change for all stimulus pairs independently of the equality of the preferences of the two options. While a computational model that represents cognitive dissonance has also been constructed ^[58]^, the model was not used for the IDM task as in the present study. It remains possible that if we build a CBL model incorporating cognitive dissonance and the model fits the behavior well, we may get different results from the present results. Furthermore, it goes beyond the integration of cognitive dissonance and the CBL model. There may be room in the future to consider the integration of RL and CBL models, which have similar formulas. This makes it possible to model complex decisions in which the EDM and IDM processes interact.

Finally, there was a possibility that preferences were formed to some extent by the first impression in IDM. Although we used novel contour shapes by following the previous study ^[9]^ to minimize the impact of initial preferential differences, we cannot rule out the possibility that value can be formed by first impression. For individuals whose preferences were formed by first impressions, it is possible that the estimated learning rate was estimated to be larger than the true value.

## Conclusion

In this study, we implemented the tasks of EDM and IDM using similar experimental procedures and applied the computational model analysis for the behavioral data of both decisions. In which EDM was followed by IDM and presented the same stimuli as EDM, we showed that the learned high values in the EDM reflect on the initial preference of the IDM. Stimuli that had been learned to have high value through EDM were also chosen in IDM, further increasing their value through IDM. The findings are consistent with structuralism, which emphasizes the impact of the external environment (e.g., through social and cultural structures) on human behavior. In contrast, from the results of the final value in IDM, stimuli that were of high value in EDM were still of high value at the end of IDM, but not as high as the novel stimuli in the most preferred IDM. These results demonstrated that externally given criteria have a strong influence on our later preferences, and at the same time, demonstrated that values formed by choice based on one’s own criteria can be higher than externally derived values (i.e., SIV). The specificity of the value learned through IDM was consistent with existentialism, which emphasizes that individuals are free to choose and establish their own identity through the choices they make. We proposed that the phenomenon of SIV in the IDM may be observed through the reward response to processing self-relevant stimuli in terms of the fundamental self-hypothesis. This study is the first to disentangle the relationship between EDM and IDM, revealing that EDM values influence IDM, and finding the phenomenon of SIV. This superiority suggests that the values learned through the EDM and IDM are likely to differ. Our findings serve as a window to the comprehensive understanding of the decision-making process.

## Methods

### Participants

38 healthy Japanese university students (male = 17, female = 21, mean age = 20.789, age-range = 18–29) participated in the experiment. All participants were native Japanese speakers, right-handed, with regular or corrected-to-normal vision. All experimental protocols were accepted by the Ethics Committee of the Graduate School of Education at the University of Hiroshima. According to the guidelines of the Research Ethics Committee of the University of Hiroshima, all participants provided informed written consent prior to participation. They were compensated for participating in the experiment.

### Stimuli

Fifteen novel contour shapes were selected from a previous study ^[59]^. We selected shapes with mild complexity (mean = 5.1, *SD* = 1.4), width (mean = 5.5, *SD* = 1.8), smoothness (mean = 4.6, *SD* = 1.7), symmetry (mean = 3.7, *SD* = 1.7), and orientation (mean = 5.0, *SD* = 2.1). The ratings of these characteristics were collected using a 9-point rating scale (1–9) ^[59]^. Additionally, these shapes had lower association values (mean = 65.71, *SD* = 6.84). The association value was the percentage (%) of respondents who gave the name of a specific object when asked to name the object recalled by the shape and those who could not write the name but said it resembled something ^[59]^. These shapes are numbered 29, 31, 35, 36, 37, 39, 42, 44, 45, 56, 63, 65, 81, 87, and 92 in the original study ^[59]^. The image used in the experiment was 800 × 600 pixels in size. Within 30 degrees of the angle of view, participants could see a picture on the screen.

In both tasks, PsychoPy ^[60]^ was used to present each pair on a white background, with one member on the left and the other on the right side of the screen. The order of the trials, as well as the presentation slides of the shapes, were randomized through the participants. The experiment was carried out on a Windows 10 PC with a 1920 × 1080 monitor.

### Task

All participants conducted EDM tasks followed by IDM (Fig 6a). Subjective preference ratings of each stimulus were conducted after the IDM.

**Fig 6.**
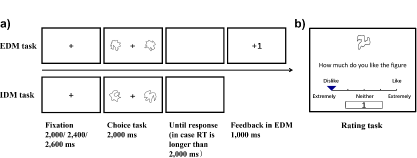
Experimental procedure. a) In the EDM task, participants were asked to choose the one considered correct from the two stimuli. In the IDM task, participants were asked to choose the one preferred from the two stimuli. Feedback was not presented in the IDM task. b) In the rating task, participants subjectively evaluated all stimuli in the IDM task on a 5-point Likert scale (1 = Extremely Dislike, 5 = Extremely Like). The rating task was conducted after the IDM task. The subjective rating data was not used for computational model analyses.

#### EDM task (Reward learning task)

Four shapes were randomly selected from 15 stimuli materials, and pairwise combinations were presented. Participants carried out six blocks of 34 trials. Each trial started with a fixing cross shown for a randomly chosen time of 2,000 ms, 2,400 ms, or 2,600 ms. Subsequently, two shapes of the fixed combinations were presented for 2,000 ms on the left and right sides of the fixation cross. Participants were asked to choose one of the two shapes considered to be correct as quickly and correctly as possible by pressing the ‘f’ key (left) or the ‘j’ key (right) on a standard computer keyboard. To limit the exposure period for each stimulus, the stimuli disappeared (i.e., turned to a white screen) after 2,000 ms. Participants could make their response even after the two shapes had disappeared, and the white screen disappeared once participants made a response. If a key was pressed within 2,000 ms, a white screen was not shown. After the response, they received correct (+1) or incorrect (0) feedback for 1,000 ms. Participants were informed that +1 indicated an increase in the number of points earned, meaning that the more points earned, the higher the reward paid to the participant after the experiment. The participants were instructed to earn as many points as possible. In each trial, each stimulus had a certain probability of receiving points after selection, and 90% vs. 10% stimuli pair or 80% vs. 20% stimuli pair appeared randomly. In addition, the left and right positions of the stimuli in each stimuli pair were also random. The participants were informed that each stimulus had a certain probability of obtaining points, but they were not informed of the specific probability. Furthermore, it was worth noting that the reward for each stimulus was generated independently in each trial. After a participant chose one stimulus of a pair and received a reward, it was difficult to infer whether another rejected stimulus could receive one. Although participants may have speculated on the possibility of the outcome of the other stimuli, the behavioral data do not support this possibility (see the supplementary materials for more details).

#### IDM task (Preference judgment task)

This task was the same as that in the previous study ^[9]^. Including the four shapes in the EDM task, all 15 shapes randomly created 105 pairs (i.e., 14 presentations per stimulus) in the IDM task, and each pair of stimuli was presented only once. There were three types of stimuli in the IDM task: novel stimuli and the stimuli presented in EDM consisted of high probability reward stimuli (90%, 80%; HP) and low probability reward stimuli (20%, 10%; LP). Participants carried out five blocks of 21 preference decision trials and were asked to choose the preferred shape among two shape stimuli presented according to their own preferential criteria in each trial. We also informed participants that there was no objectively correct answer in this task. Stimuli were presented in the same manner as the EDM task, except there was no feedback after the choice.

#### Rating task

We conducted a subjective rating task for each shape stimuli (Fig 6b) to examine the relationship between subjective preferences and final stimulus values estimated by computational model analysis for IDM behavioral data. Following the IDM task, participants carried out a subjective preference rating task. In the rating task, participants were asked to determine their subjective preference, graded on a 5-point Likert scale (1 = Extremely Dislike, 5 = Extremely Like) for each shape. It is worth noting that for the IDM task, we did not use the experimental paradigm with subjective ratings, such as the free choice paradigm ^[19]^, or rate-rate-choice ^[61]^, to avoid the influence of noise-contaminated subjective ratings on CIPC measurement ^[62]^. As a result, preference rating data in the Likert scale was not included in the computational model analyses.

### Classical analysis of the behavioral data

We first confirmed the correct response rate for each stimulus pair in the EDM task to gauge whether participants learned through the EDM task.

To investigate whether the value learned in EDM had an effect on IDM, prior to computational model analyses, we compared the chosen frequency of the three types of stimuli (HP, LP, and novel) in the IDM task using the paired *t*-test with Holm multiple-comparison correction. The chosen frequency of each type of stimuli was calculated by dividing the number of times it was chosen across all trials by the number of times it was presented.

### Computational models

To investigate how the values learned in EDM reflected IDM, we prepared four CBL models with different initial stimulus values (Table 1), which established by the previous study ^[9]^. The CBL model’s learning process involves increasing the value of chosen items while decreasing the value of rejected items. The CBL model is written as follows:

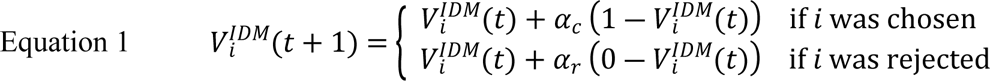

The values (*V*^*IDM*^) in CBL models were updated based on whether a participant chose or rejected it. The updated *V*^*IDM*^ was kept constant until the trial in which the stimulus was presented. The degree of value change followed by choice was determined by the learning rate (*α*_*c*_ or *α*_*r*_). When item *i* was chosen, the learning rate (*α*_*c*_) was multiplied by 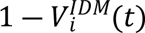 and added to the value at trial *t*, as if it were the prediction error for a correct response (feedback is +1) in the RL model (see the supplementary materials Equation s1). In case item *i* was rejected, the learning rate (*α*_*r*_) was multiplied by 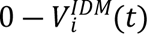 and added to the value at trial *t* as if it were the prediction error in the RL model for an incorrect response (feedback is 0) in the RL model.

The typical RL model in Equation s1 (see the supplementary materials) does not update the values of the rejected items. In contrast, in CBL, participants updated the value of items based on their own choices; hence, rejected items were considered incorrect answers, and their values were updated to decrease. Notably, although the EDM and IDM are similar in updating values through differences in existing values and feedback, there are differences in updating the values of both or chosen options. These differences arise from task design. In a typical EDM task, including our EDM task, the feedback for the two items is independently determined by probability and participants are only informed of the feedback of the chosen option. Therefore, knowing whether the rejected item was the correct answer was difficult, and only the value of the chosen option would increase or decrease based on feedback from external circumstances ^[63]^. In contrast, in the IDM task, it is clear that what they choose is preferred and what they do not choose is not preferred. Thus, the value of the chosen option increases and the value of the rejected option decreases ^[19]^. Zhu et al. ^[9]^ compared CBL models that update the value of both chosen and rejected models that update only one of them and reported that models that update both values have a better fit to the behavior. We used the same IDM task as Zhu et al. ^[9]^. Although not directly related to the aim of this study, we confirmed that the behavioral data from this study are better suited for an RL model that only updates chosen options rather than one that updates both chosen and rejected options (see the supplementary materials).

The initial values of the stimulus types (novel, HP, or LP), wherein the models differed, were estimated as free parameters. Model 1 represented no influence of the values learned in EDM, and the initial value *η* (0 ≦ *η* ≦ 1) was a free parameter, which was the same for all stimulus types in the IDM task. Model 2 represented that only the initial value of the HP stimuli can reflect the EDM value and differed from the other stimuli in the IDM task. The initial value of HP stimuli (*η*_*HP*_) was a free parameter, whereas the initial value of the LP stimuli was fixed at 0.5, the same as with novel stimuli. In contrast to Model 2, in Model 3, only LP was a free parameter, and the others were fixed at 0.5. In Model 4, both HP and LP were free parameters, and the novel stimuli were fixed at 0.5. Since the previous study ^[9]^ reported that the model that used different learning rates for chosen and rejected items was unsuitable for model comparison, the above four models were created based on the model that used the same learning rates (i.e., *α*_*c*_ = *α*_*r*_) as the main model-based analysis. Model 1 is a null model that assumes no effect of the EDM on the IDM. If the fit of the other models is superior to that of Model 1, this indicates an effect of the EDM on the IDM. Because behavioral data are more likely to fit a model with a higher number of parameters ^[64]^ and to avoid the issue of parameter setting for the initial values of other models affecting the fit of Model 1, we set the initial values of Model 1 as free parameters under the constraint of no influence from the EDM. Thus, if other models provided a better fit, we could clearly determine the effect of the EDM on the IDM.

To calculate the probability of choice in the CBL models, the softmax function was applied to the value difference between the two options.

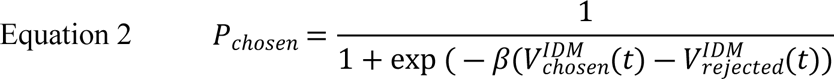

In trial *t*, 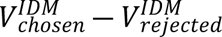 with parameter *β* was used to determine the value of *P*_*chosen*_, which was used to represent the probability that the model chooses the option the participant chose. *β* determined the softmax function’s slope. The higher the value, the more the decision was based on the values, while the lower the value, the more random the decision was and the less reliant on the value.

### Simulations and model-based behavioral data analyses

All CBL models (Table 1) underwent both simulations of parameter and model recoveries. CBL models that passed the parameter recovery performed model recovery, testing whether the model that generated the artificial data best fit the same model. Finally, we fitted the actual behavioral data to the CBL models to determine which model best explained the behavioral data.

All subsequent simulations and actual data analyses were performed on R ^[65]^. Hierarchical Bayesian method was used to derive model parameters, and the calculation process was completed by rstan package ^[66]^. This method assumes that a common distribution within the group generates each participant’s parameters (e.g., *α*_*n*_, *β*_*n*_). As shown in the following Equation 3 and Equation 4, the parameters *α* and *β* of participant *n* were assumed to be generated from the normal distributions of *μ* and *σ*^2^. As with the prior distributions of *μ* and *σ*^2^, we used the uniform distribution. At the same time, we truncated the normal distribution to ensure that the generated parameters were within a certain range.

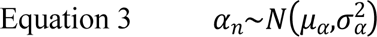

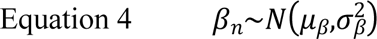

The parameters at the population level (*μ*_*α*_, 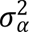, *μ*_*β*_, 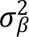) were used as free parameters to infer from data. At the population level, the distribution parameters took a prior distribution into account, and the distribution calculated a posterior distribution using the Bayes estimator. The posterior distribution of parameters was obtained by the Markov Chain Monte Carlo method (MCMC).

### Simulation 1 (Parameter recovery)

We conducted parameter recovery simulations for the CBL models to evaluate whether the experimental settings and models met the goal of estimating model parameters from behavioral data ^[67]^. We tested whether model parameters used to produce artificial behavioral data could be estimated by model fitting to the artificial data. As parameter recovery indices, Pearson’s correlation coefficient was calculated between the simulated and fitted parameters.

We used the same settings as the actual experimental design when generating the artificial behavioral dataset. In each model, we generated artificial data with 15 stimuli and 105 trials for 38 people. Initial values for stimuli were set as in Table 1. The *α* (0 ≦ *α* ≦ 1), *β* (0 ≦ *β* ≦ 20), and *η* (0 ≦ *η* ≦ 1) were generated from the normal distribution of *μ* and *σ*^2^. *μ*_*α*_, 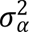, *μ*_*η*_, and 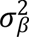 were generated from the uniform distribution ranging from 0 to 1, while *μ*_*β*_ and 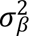 were generated from the uniform distribution ranging from 0 to 20 and 0 to 10, respectively.

### Simulation 2 (Model recovery)

Model recovery was conducted to test whether the true model showed the best fit for the data generated by that model under the experimental design. The widely applicable Bayesian information criterion (WBIC) ^[64]^ was used to assess the relative goodness of fit of the models. As shown in Equation 5, the-WBIC is equal to the approximate value of log marginal likelihood.

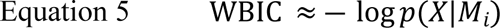

*p*(*X*|*M*_*i*_) is the marginal likelihood, which is the probability of generating data X given by the model *M*_*i*_.

More specifically, we first used the MCMC method after performing the transformation in Equation 6 to estimate the log posterior density 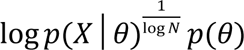.

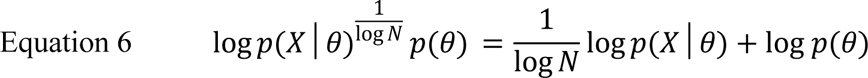

*θ* was the vector of parameters such as *α*, *β*. *N* represented the sum of all trials across participants.

The WBIC was then calculated from the S samples of MCMC as Equation 7.

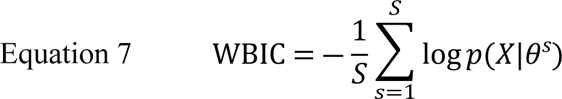

S for model recovery was 5000 samples, while those for parameter recovery and behavioral data analysis were 10000.

A smaller WBIC indicated a better fit of the model to the data. The four individual models shown in Table 1 generated artificial behavioral data in the same way with Simulation 1. Subsequently, each data set was fitted to all models and judged which model best fit the data using WBIC.

To compare which model had the higher probability of generating data, we calculated the Bayes factor (BF). The BF was calculated by the ratio of the marginal likelihood of the two models and was calculated using the marginal likelihood of the model used to generate the data as the numerator. A previous study ^[68]^ referred to evaluated Bayes factor of 1–3 as not worth mentioning, 3–20 as positive, 20–150 as strong, and more than 150 as very strong.

### Models fit to the behavioral data

To conduct computational model analysis of the actual behavioral data in the IDM task, we first applied participants’ behavioral data to all CBL models passing parameter recovery. As in Simulation 2 (Model recovery), WBIC was calculated as an index of model fit, and BF was used for inter-model comparison. The estimated model parameters and values of each stimulus from the best fit model were used for further analyses.

### Additional model comparison between the CBL and the models without value updates in IDM

The CBL models used in this study were based on those developed by Zhu et al. ^[9]^. Although in their study, the CBL models were validated to show a change in the value of chosen and rejected options after preference-based choices, we cannot rule out the possibility that the value of the stimuli in this study remained unchanged after choices. Therefore, an additional computational model analysis was performed to investigate whether the value did not change in the IDM. Additional models without value changes were constructed based on the best-fit CBL model for the behavioral data in Models 1–4. Specifically, we created four additional models (Table 4): Model A, in which only the value of the HP stimuli was not updated; Model B, in which both the HP and LP stimuli were not updated and only the value of the novel stimuli was updated; Model C, in which only the value of the novel stimuli was not updated; and Model D, in which the values of all the stimuli were not updated. All new models were analyzed through simulations and the fitting of behavioral data.

## Data availability

The data of each participant (correct response rate, chosen frequency, subjective evaluation, and the value of each stimulus estimated by the computational modeling analysis), and the code used for the analyses are available from: https://doi.org/10.6084/m9.figshare.20400666.v3

## Acknowledgments

JST COI Grant Number JPMJCE1311 and JSPS KAKENHI Grants 18K03177, 18K03173, 22K07328, and 22H01083 supported this research.

## Author contributions statement

J.Z. designed the experiments, analyzed the data, and wrote the manuscript. All the authors reviewed and revised the manuscript.

## Competing interests

The authors declare no competing interests.

## Notes

### Competing Interest Statement

The authors have declared no competing interest.

